# CometChip Enables Parallel Analysis of Multiple DNA Repair Activities

**DOI:** 10.1101/2021.01.19.427336

**Authors:** Jing Ge, Le P. Ngo, Simran Kaushal, Ian J. Tay, Elina Thadhani, Jennifer E. Kay, Patrizia Mazzucato, Danielle N. Chow, Jessica L. Fessler, David M. Weingeist, Robert W. Sobol, Leona D. Samson, Scott R. Floyd, Bevin P. Engelward

## Abstract

DNA damage can be cytotoxic and mutagenic and is directly linked to aging, cancer, and heritable diseases. To counteract the deleterious effects of DNA damage, cells have evolved highly conserved DNA repair pathways. Many commonly used DNA repair assays are relatively low throughput and are limited to analysis of one protein or one pathway. Here, we have explored the capacity of the CometChip platform for parallel analysis of multiple DNA repair activities. Taking advantage of the versatility of the traditional comet assay and leveraging micropatterning techniques, the CometChip platform offers increased throughput and sensitivity compared to the traditional comet assay. By exposing cells to DNA damaging agents that create substrates of Base Excision Repair, Nucleotide Excision Repair, and Non-Homologous End Joining, we show that the CometChip is an effective method for assessing repair deficiencies in all three pathways. With these advanced applications of the CometChip platform, we expand the efficacy of the comet assay for precise, high-throughput, parallel analysis of multiple DNA repair activities.

## INTRODUCTION

DNA damage promotes cancer, aging, neurological disorders and heritable diseases. Exposure to genotoxins is unavoidable, as DNA damaging agents are ubiquitous both in our environment and within our cells. To prevent adverse effects associated with damage to the genome, all species have evolved defenses for removing and repairing damaged DNA (Chatterjee and Walker 2017; Errol et al. 2006). Over 200 human proteins have been categorized as *bona fide* DNA repair proteins (Wood et al. 2001; Wood et al. 2005, 2020) (12 January 2021, date last accessed). Most fall into the five classical DNA Repair pathways of Base Excision Repair (BER), Mismatch Repair (MMR), Nucleotide Excision Repair (NER), Homologous Recombination (HR) or Non-Homologous End-Joining (NHEJ). It is now well established that deficiencies in some DNA repair proteins promote genomic instability and can lead to carcinogenesis (Alberts 2009; Andre et al. 2020; Chatterjee and Walker 2017; Ciccia and Elledge 2010; Errol et al. 2006; Jackson and Bartek 2009; Latham et al. 2019; Lord and Ashworth 2012; Torgovnick and Schumacher 2015). Importantly, DNA damaging agents are also frequently used as chemotherapeutics to target rapidly dividing tumor cells (Butler et al. 2018; Lord and Ashworth 2012; Torgovnick and Schumacher 2015). Therefore, understanding DNA repair capacity and the coordination of DNA repair pathways is important not only in revealing disease etiology, but also in predicting chemotherapeutic efficacy (Andre et al. 2020).

A wide variety of methods are used to measure DNA damage and repair, but we lack tools to evaluate damage induction and repair in multiple DNA repair pathways in parallel (Valdiglesias et al. 2011). For example, whereas DNA sequencing can identify DNA repair gene variants and rare mutations, the consequences are not always clear (Lo et al. 2003). Transcriptional profiling is often used to evaluate expression of DNA repair genes, however transcript levels do not always reflect DNA repair capacity (Nagel et al. 2014a). Direct activity assays using cell lysates are highly effective and have been used in population studies (Edwards et al. 2015; Paz-Elizur et al. 2003; Paz-Elizur et al. 2006; Paz-Elizur et al. 2020; Somuncu et al. 2020), however they are not always conducive to analysis of multiple pathways in parallel (Geng et al. 2011; Georgiadis et al. 2012; Leitner-Dagan et al. 2012; Redaelli et al. 1998; Svilar et al. 2012; Zhong et al. 2002). To enable analysis of multiple DNA repair pathways, a fluorescence-based multiplex flow-cytometric host cell reactivation assay (FM-HCR) was recently developed (Nagel et al. 2014b). Here, we describe an alternative approach, which is to measure the activity of multiple DNA repair pathways using the single cell electrophoresis assay, known as the comet assay.

The comet assay can be used to measure the formation and repair of strand breaks and alkali labile sites, which arise from exposure to DNA damaging agents or from endogenous cellular processes. This assay was first developed by Ostling, Johansen, and Singh more than 30 years ago (Ostling and Johanson 1984; Singh et al. 1988), and it has been improved by many laboratories (Azqueta et al. 2014; Hartmann et al. 2003; Langie et al. 2006; Olive and Banath 2006). The comet assay is based on the principle that, in an agarose gel under an electrophoretic field, damaged DNA migrates farther than undamaged DNA. In this assay, cells are exposed to test conditions, embedded in agarose, subjected to electrophoresis, and stained with a nucleic acid dye. DNA damage can also be directly quantified by analyzing the extent of DNA migration. Multiple versions of the comet assay have been developed to measure repair of different kinds of damage in the genomes of live cells (A Collins et al. 1997; Collins 2004). While the traditional comet assay has gained increasing popularity for studies of DNA damage and repair (Anderson et al. 2019; Bankoglu et al. 2019; Collins et al. 2019; Gajski et al. 2020a; Gajski et al. 2020b; Hobbs et al. 2020; Kruszewski et al. 2019; Misik et al. 2019; Vodicka et al. 2019; Witte et al. 2007), it has not been widely adopted for multi-pathway analysis, in part due to low throughput, high inter-experimental and inter-laboratory variability, and the requirement for laborious analysis (Azqueta et al. 2011; Azqueta et al. 2020; Collins et al. 2014; Ersson et al. 2013; Forchhammer et al. 2008; Forchhammer et al. 2010; Forchhammer et al. 2012; Moller et al. 2004).

We have previously exploited microfabrication technologies to develop the CometChip, a mammalian cell array format that increases comet assay throughput and reproducibility (Figure 1) (Weingeist et al. 2013; Wood et al. 2010). The technology uses photolithography to fabricate PDMS molds that have arrayed microposts to cast microwells in agarose (Figure 1A). Each microwell can be tuned to the size of a single cell. Cells are then loaded in the microwells by gravity and encapsulated with low melting point agarose (Figure 1B). By patterning cells into arrays, the technology minimizes the area required per sample, prevents overlapping comets, and places comets on a shared focal plane, such that only 1 or 2 images are required, rather than hundreds of images as is currently the standard for the traditional comet assay. Comets are then analyzed using freely available in-house software (Ge et al. 2015; Weingeist et al. 2013; Wood et al. 2010). Whereas in the traditional comet assay each sample requires its own slide, the CometChip fits each sample into a single well of a standard 96-well (Figure 1). Previous studies show that, when compared to the traditional comet assay, the CometChip platform increases throughput by more than two orders of magnitude, and reduces noise (Ge et al. 2015; Weingeist et al. 2013). The CometChip, therefore, potentiates sensitive measurement of DNA damage and repair in live cells in a high throughput fashion.

**Figure 1.**
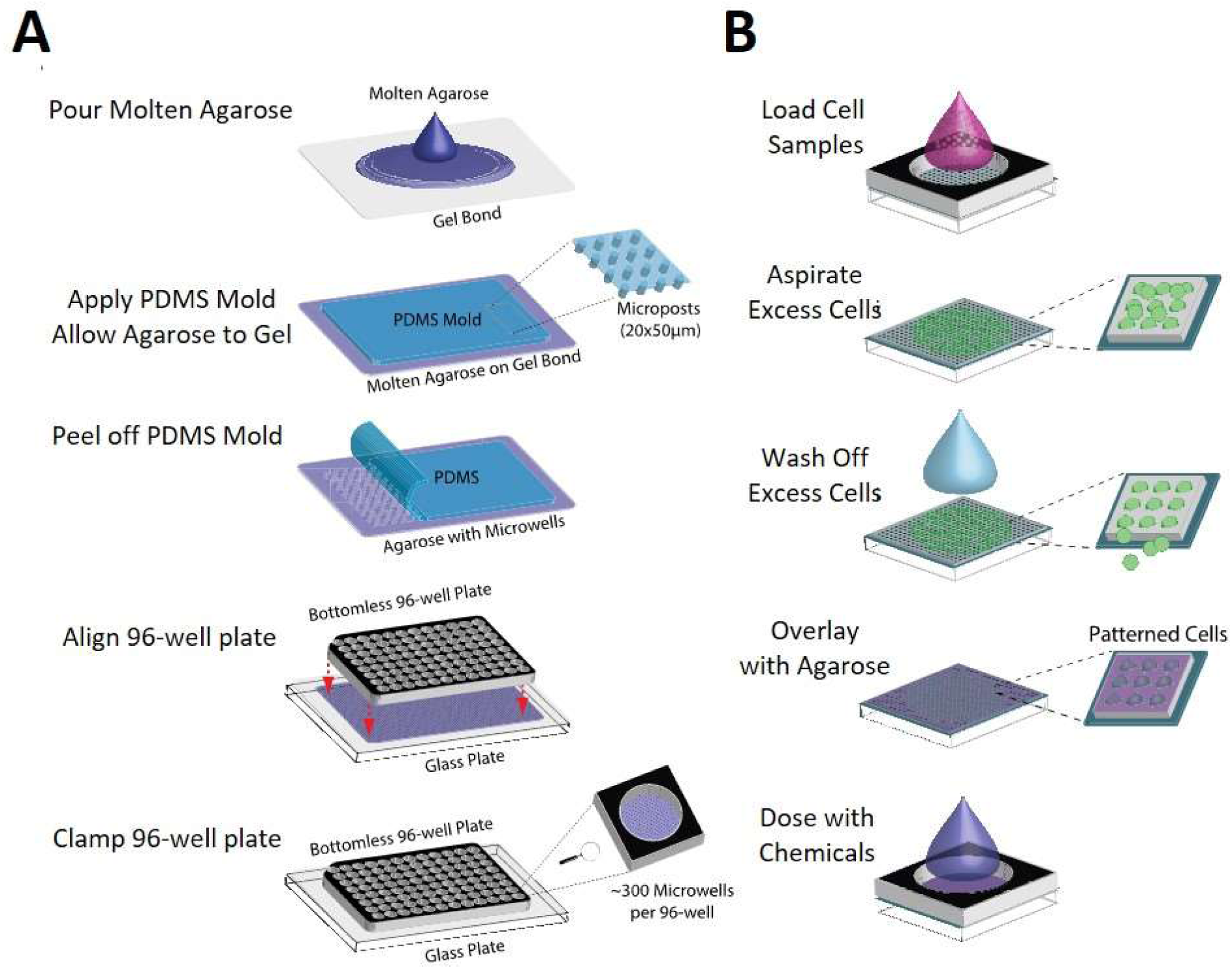
Patterning cells in the 96-well CometChip (A) Procedures of making the 96-well CometChip using microfabricated PDMS mold. (B) Zoomed in view of cell patterning procedures in one well of the 96-well CometChip.

DNA damage and repair responses to genotoxic stress involve a complex network of multiple proteins and pathways. Two main pathways for excision of damaged DNA nucleotides are BER (for single base lesions) and NER (for bulky lesions and intrastrand DNA crosslinks). Both pathways have strong relevance to human health. People with BER gene polymorphisms and deficiencies have been shown to have increased cancer susceptibility (AlMutairi et al. 2015; Chaim and Nagel 2017; Chatterjee and Walker 2017; Cheadle and Sampson 2007; Goode et al. 2002; Hakem 2008; Leitner-Dagan et al. 2012; Leitner-Dagan et al. 2014; Li et al. 2014; Nagel 2017; Sevilya et al. 2014; Sevilya et al. 2015; Sliwinski et al. 2007; Sweasy et al. 2006) and show a link to degenerative diseases (Allocca et al. 2017; Ebrahimkhani et al. 2014; Kisby et al. 2011; Meira et al. 2009). Similarly, reduced NER capacity is also associated with increased risk of cancer (Azqueta et al. 2014; Berndt et al. 2006; Chang et al. 2008; Dai et al. 2019; He et al. 2016; He et al. 2014; Huang et al. 2006; Li et al. 2020; Slyskova et al. 2011; Su et al. 2019; Thoms et al. 2007; Tian et al. 2020; Yan et al. 2020; Zhu et al. 2018). DNA double strand breaks (DSBs) are repaired primarily by NHEJ and HR. In the case of NHEJ, polymorphisms in essential genes predispose individuals to breast, lung, brain, and liver cancer (Errol et al. 2006; Sishc and Davis 2017). Taken together, DNA repair is increasingly recognized as being critical for suppressing cancer, making a multi-pathway analysis platform relevant for learning more about DNA repair, performing epidemiological studies, and personalized medicine.

In order to leverage the CometChip for broad assessment of DNA repair capacity, we set out to analyze its utility for assessing repair of lesions from DNA damaging agents that create substrates for three major DNA repair pathways: BER, NER, and NHEJ. Using genetically defined cell lines depleted for key DNA repair enzymes, we evaluated the potential efficacy of the comet assay for assessment of these repair deficiencies using model DNA damaging agents. In order to study multiple DNA repair pathways, we assessed multiple time points using cell lines with varied DNA repair capacity in triplicate for each experiment and performed each experiment three times. Altogether, over 1,500 samples were analyzed (equivalent to 150,000 comets), which is far greater than what is feasible using the traditional comet assay, both because of the requisite labor, and also because of noise from sample to sample and from experiment to experiment. Our results demonstrate that the CometChip enables detection of a variety of DNA lesions and support its use for high throughput applications that require assessment of multiple cell types, chemical conditions, and repair time points.

## MATERIAL AND METHODS

### Cells and cell culture

TK6 human lymphoblastoid cells were a gift from Dr. William Thilly. TK6 and MCL-5 cells (Crespi et al. 1991) were cultured in suspension in 1x RPMI 1640 medium with L-glutamine (Invitrogen) supplemented with 10% horse serum (Invitrogen) and 100 units/ml penicillin-streptomycin (Invitrogen) (Liber and Thilly 1982; Skopek et al. 1978). Blood from one anonymous healthy donor collected in a sodium heparin Vacutainer collection tube was purchased from Research Blood Components, Brighton, MA. Primary lymphocytes (peripheral blood mononuclear cells, PBMCs) were isolated from the blood using the standard Ficoll gradient density centrifugation. The PBMC pellet was suspended in freezing medium (40% RPMI-1640 + 50% HI-FBS + 10% DMSO). Vials were stored at-80°C. Cryopreserved lymphocytes were rapidly thawed at 37°C and resuspended in stimulation medium (RMPI-1640 + 20 % HI-FBS + 100 U/mL Pen-Strep + 5 μg/mL PHA-L) and T-lymphocytes were stimulated for 3 days at 37°C and 5% CO_2_. Wildtype, AagTg, Aag^−/−^, and *Polβ*^−/−^mouse embryonic fibroblast (MEF) cell lines were generated as described in (Sobol et al. 2003). XPCS1RO and XPCS1RO + XPG human fibroblasts were a gift from Dr. Orlando D Schärer (Chemical&Cancer Biology Branch, Institute of Basic Science – Center for Genomic Integrity, Ulsan, South Korea (Ellison et al. 1998)). MEFs and human fibroblasts were cultured in DMEM (Invitrogen) supplemented with 10% FBS (Atlanta Biologicals, Atlanta, GA) and 100 units/ml penicillin-streptomycin (Invitrogen). Human glioblastoma cell lines M059J and M059K (from American Type Culture Collection) were cultured in DMEM/F12 medium (Invitrogen) supplemented with 10% FBS (Atlanta Biologicals, Atlanta, GA), MEM Non-Essential Amino Acids (Invitrogen) and 100 units/ml penicillin-streptomycin (Invitrogen). LN428 glioblastoma cell lines were described previously (Fang et al. 2014).

### CometChip Fabrication

Figure 1A illustrates the 96-well CometChip fabrication using a PDMS mold. Briefly, 15 mL of molten normal melting point agarose was applied to GelBond® (Lonza, Switzerland) in a rectangular one-well plate (VWR). The agarose was allowed to gel with the PDMS mold on top. Phosphate buffered saline (PBS) was added to aid in removal of the mold, which revealed patterned microwells across the surface of the gel. The gel was sandwiched between a glass plate and a bottomless 96-well plate and the setup was secured with 2’’ binder clips to create the 96-well CometChip. At least 10,000 cells were added to each macrowell and allowed to settle by gravity in complete growth media at 37°C, 5% CO_2_. Excess cells were washed off after 15-30 min and the bottomless plate was removed in order to enclose the arrayed cells in a layer of 1% low melting point agarose (Figure 1B).

### Chemical Treatments

Hydrogen peroxide (H_2_O_2_) (Sigma) was diluted in cold PBS to the desired dose immediately before use. Encapsulated cells were immersed in ice cold H_2_O_2_ and incubated for 20 minutes prior to lysis. Methyl methanesulfonate (MMS) (Sigma) was diluted in PBS and cells were treated for 30 min at 37°C. Negative control cells were treated with complete media with the corresponding DMSO concentrations. After treatment, cells were either replenished with fresh media to allow measurement of repair kinetics or trypsinized and resuspended in media to load onto the CometChip for dose response experiments. After loading and encapsulation, gels were immediately lysed and alkaline comet assay was performed. For dead cell comet analysis, cells embedded in the CometChip were lysed for 4 hours at 4°C, washed with 1x PBS, and then exposed to H_2_O_2_.

### Enzyme Treatments

Following chemical treatment with H_2_O_2_ or MMS, encapsulated cells were lysed overnight. H_2_O_2_-treated lysed cells were incubated with 0.8 units/mL Fpg enzyme (New England Biolabs) in Fpg reaction buffer (40mM HEPES, 0.1M KCl, 0.5mM EDTA, 0.2mgl/mL BSA, pH 8.0) for 20 min at 37°C. MMS-treated cells were lysed overnight, washed in hAAG enzyme reaction buffer (20 mM Tris-HCl, 10mM (NH_4_)_2_SO_4_, 10mM KCl, 2 mM MgSO_4_, 0.1% Triton®X-10, pH 8.8 at 25°C) three times for 15 min, incubated with 25 units/mL hAAG enzyme (New England Biolabs) at 37°C for 15 min, and washed with PBS before unwinding. For controls, lysed cells were incubated with reaction buffer or with PBS under the same conditions.

### UV Irradiation and Repair

TK6 cells were harvested and resuspended in culture medium supplemented with 10 mM L-glutathione (L-GSH) (Sigma). Following loading in the CometChip encapsulation with 1% low melting point agarose, gels were cut into 3×3 macrowell sizes and individually exposed to UV light using a 254 nm UVC lamp (UVG-11, UVP, Upland, CA) calibrated with a UVX radiometer (UVP, Upland, CA) to 0.5 ± 0.1 J/m^2^/s. Exposure was performed in the dark at 4°C and cells were transferred to culture media with 10 mM L-GSH to allow for repair at 37°C. Cell lysis and alkaline comet assay were performed subsequently.

### Ionizing Radiation Exposure

Cells were first loaded into the CometChip and overlaid with low melting point agarose. The gels were then cut and exposed in smaller pieces. Irradiation was performed using Gammacell 220E (MDS Nordion, Canada), which delivers ionizing radiation (IR) from a Co^60^ source at an approximate rate of 100 Gy/min (depending on source age). Cells were irradiated at room temperature. To evaluate repair kinetics, wells were allowed to repair damage in media for varying time intervals, after which each well was filled with cell lysis buffer to terminate repair.

### Alkaline Comet Assay

The alkaline comet assay was performed as described previously (Ge et al. 2014). Briefly, for dose-response experiments cells embedded in agarose gels were lysed in lysis buffer (10 mM Tris-HCl, 100 mM Na_2_EDTA, 2.5 M NaCl, pH 10 with 1% Triton X-100 added 20 min before use) overnight at 4°C. After lysis, the gels were transferred to an electrophoresis chamber filled with alkaline unwinding buffer (0.3 M NaOH and 1 mM Na_2_EDTA, pH>13) for 40 min at 4°C. Electrophoresis was conducted with the same buffer at 4°C for 30 min at 1 V/cm and 300 mA. Percent tail DNA was used to quantify DNA damage for alkaline comet assay.

### Neutral Comet Assay

The neutral comet assay was performed using a modified version of the neutral comet protocol as previously described (Ge et al. 2014). Briefly, following treatments with DNA damaging agents, cells embedded in agarose gels were lysed for 4 hours at 43°C (2.5 M NaCl, 100 mM Na_2_EDTA, 10 mM Tris, 1% N-Laurylsarcosine, pH 9.5 with 0.5% Triton X-100 and 10% DMSO added 20 min before use). For evaluation of repair kinetics after treatments with DNA damaging agents, the cells were lysed by adding lysis buffer to the appropriate wells of the multiwell plate at 37°C while the repair kinetics time course was carried out and then transferred to 43°C overnight to complete the lysis step. After lysis, the gels were washed three times for 30 min with the electrophoresis buffer (90 mM Tris, 90 mM Boric Acid, 2 mM Na_2_EDTA, pH 8.5). Electrophoresis was conducted at 4°C for 1 hour at 0.6 V/cm and 6 mA. Tail length was used to quantify DNA damage for neutral comet assay.

### Fluorescence Imaging and Comet Analysis

After electrophoresis, agarose gels were neutralized in 0.4 M Tris, pH 7.5 neutralizing buffer twice for 15 min and then stained with 1x SYBR Gold diluted in PBS (Invitrogen). Images were captured at 40x magnification using an epifluorescence microscope (Nikon Eclipse 80i, Nikon Instruments, Inc., Melville, NY, USA) with a 480 nm excitation filter. Comet images were automatically analyzed using custom software written in MATLAB (The Mathworks, Inc., Natick, MA, USA) (Wood et al. 2010). Outputs from Guicometanalyzer were processed by Comet2Excel, an in-house software developed in Python (Python Software Foundation, Python version 2.7.10). All software is freely available upon request. At least 100 comets were analyzed per condition.

## RESULTS

### Detecting BER activity

The basis for the comet assay is that during electrophoresis, damaged DNA migrates through a matrix more readily than undamaged DNA. DNA is normally highly supercoiled, and thus recalcitrant to DNA migration. When strand breaks are introduced by exposure to a DNA damaging agent, there is a loss of superhelical tension, enabling migration of long loops of DNA and fragmented DNA to create what looks like a comet (Figure 2A) (Ostling and Johanson 1984). The comet assay detects both single strand breaks (SSBs) and double strand breaks (DSBs); however, SSBs predominate in the cell under most conditions and as a result, the detection of DSBs is usually negligible under the alkaline conditions used for SSB detection. The assay also detects abasic sites and alkali sensitive moieties that are converted to SSBs under alkaline conditions (Collins 2004; Hartmann et al. 2003; Olive and Banath 2006; Ostling and Johanson 1984; Singh et al. 1988). Importantly, base modifications caused by exposure to reactive oxygen species or alkylating agents are often not directly detectable by the assay due to their stability under alkaline conditions (Figure 2B), which is consistent with the fact that exposure to an oxidizing agent does not produce detectable DNA damage in dead cells (Figure 2C). However, in live cells, BER enzymes quickly recognize and excise oxidized and alkylated bases, creating BER intermediates that can be detected by the comet assay (Figure 2B and C). Thus, while damaged bases generally cannot be directly detected using the comet assay, downstream BER intermediates (e.g., abasic sites and SSBs) are detectable, enabling studies of BER activity.

**Figure 2.**
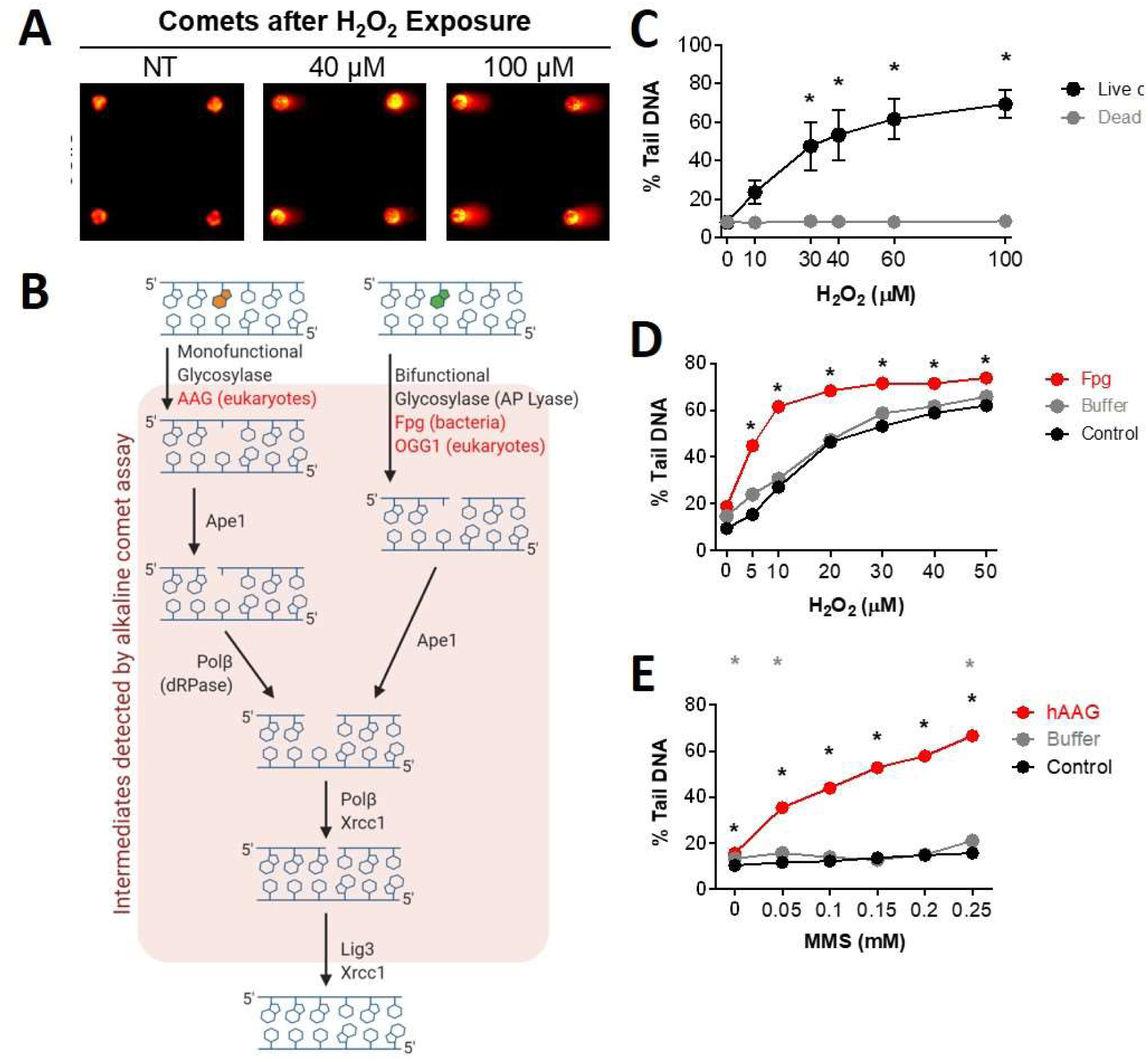
Measuring DNA BER of oxidative lesions. (A) Representative arrayed comet images. (B) Schematic of the mammalian BER pathway. H_2_O_2_ dose response curves when exposed to multiple doses of H_2_O_2_ for 20 min at 4°C for (C) live and lysed (dead) TK6 lymphoblasts and (D) MCL-5 cells without Fpg, with Fpg buffer, and with Fpg enzyme + Fpg buffer. (E) MMS dose response curves when exposed to multiple doses of MMS at 37°C for 30 min of MCL-5 cells without hAAG, with hAAG buffer, and with hAAG enzyme + hAAG buffer. *p < 0.05, Student’s t-test (one-tailed, paired), comparing to WT at each time point (black stars show hAAG compared to WT, grey stars show Buffer only compared to WT).

Although damaged DNA bases do not affect DNA migration and thus cannot be detected using the comet assay, conditions that promote DNA cleavage at sites of unrepaired DNA base lesions is an effective means of unveiling their presence. To accomplish this, DNA harboring damaged bases is incubated with a bifunctional DNA glycosylase that not only recognizes and removes the damaged base, but also cleaves the backbone to introduce a nick. To use this approach for oxidative damage, cells can be treated with H_2_O_2_, lysed, rinsed and then incubated with purified formamidopyrimidine DNA glycosylase (Fpg) from *E. coli*, which removes oxidized purines such as 8-oxoguanine (8oxoG) and for which the lyase activity introduces a nick. In essence, Fpg converts undetectable 8oxoG lesions into detectable strand breaks. This is a commonly used approach (Azqueta et al. 2013; Azqueta and Collins 2013; Collins 2004; Collins et al. 2014; Gedik et al. 2005) and we have demonstrated here that this is also an effective approach for MCL-5 lymphoblastoid cells using the CometChip (Figure 2D).

We demonstrate that this approach can be applied to reveal the presence of alkylated DNA bases as well. Cells were exposed to the alkylating agent MMS, which produces DNA alkylation lesions such as 3-methyladenine (3MeA) and 7-methylguanine (7MeG). After exposure, cells were lysed and incubated with the purified human BER glycosylase hAAG, which excises alkylated DNA bases, and the comet assay was performed. Although hAAG is a monofunctional glycosylase that removes damaged bases but does not cleave the backbone, we were nevertheless able to detect AAG-induced strand breaks, because ring-opened abasic sites are readily converted into strand breaks under alkaline conditions (Hartwig et al. 1996; Mattes et al. 1986; Prakash and Gibson 1992). Analogous to the Fpg assay, incubation with purified hAAG produced strand breaks detectable by comet assay (Figure 2E). Thus, we expand upon the approach of using a purified BER glycosylase to reveal the presence of an additional class of DNA lesions.

To more closely examine the role of BER proteins in the repair of oxidative and alkylation damage using the CometChip, we used knockout mouse embryonic fibroblasts (MEFs). Specifically, we used the CometChip to assess the repair capacities of MEFs with deficiencies in BER proteins. For oxidative damage, we compared the levels of BER intermediates in wild-type, *Ogg1* null, and *Polβ* null cells. OGG1 is the main glycosylase responsible for removing oxidized purines (e.g., 8oxoG) and Polβ is a key polymerase involved in gap filling for short-patch BER (Figure 2B). For *Polβ*^−/−^ cells, we observed more DNA damage after exposure to H_2_O_2_ (Figure 3A). These results indicate that BER intermediates are more persistent when Polβ is absent, which is consistent with previous biochemical studies showing that Polβ is rate-limiting (Srivastava et al. 1998). We also observed suppression of detectable DNA damage in *Ogg1*^−/−^ cells exposed to H_2_O_2_ (Figure 3B). This result is indicative of reduced initiation of the BER pathway. As such, there is reduced conversion of undetectable base lesions into detectable BER intermediates.

**Figure 3.**
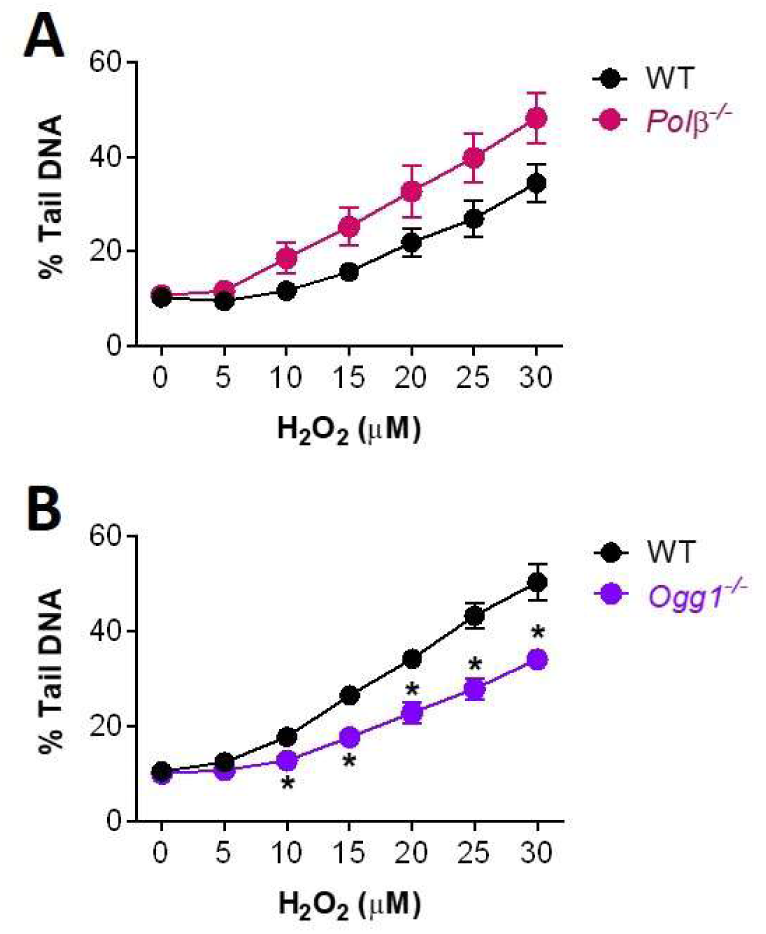
Measuring DNA repair in the absence of BER proteins. (A) and (B) Repair kinetics of mouse embryonic fibroblasts after treatment with 50 μM H2O2 at 4°C for 20 min. All NT cells were exposed to 1x PBS as control. Repair was performed in media at 37°C. Data points and error bars represent averages and standard errors of the mean (SEM), respectively, of three independent experiments (n ≥ 3) with at least 100 comets scored for each condition in each experiment. *p < 0.05, Student’s t-test (one-tailed, paired), comparing to WT at each time point.

### Detection of DNA repair using the CometChip

To learn more about the efficacy of the comet assay for studies of BER enzymes involved in repair of alkylation damage, we analyzed the impact of AAG and Polβ on repair kinetics for MMS-induced base lesions. After incubation with MMS for 30 min at 37°C, we see significant induction of DNA damage in wildtype (WT) cells (Figure 4A). However, we do not see rapid completion of the BER pathway for MMS-induced DNA damage (Figure 4B), possibly because MMS is more stable than H_2_O_2_, and thus MMS might continue to induce new damage throughout the exposure period.

**Figure 4.**
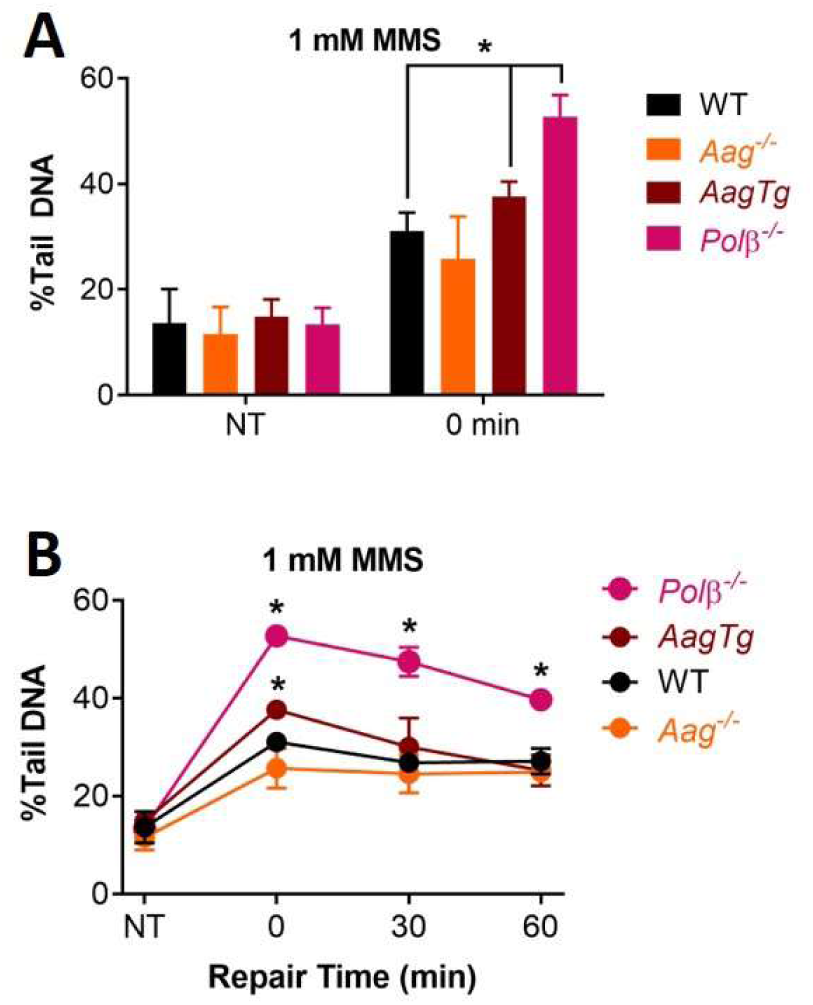
Measuring DNA BER of alkylation damage. (A) Analysis of the damage levels for non-treated (NT) and 0 min data for mouse embryonic fibroblasts after treatment with 1mM MMS at 37°C for 30 min. (B) Repair kinetics of mouse embryonic fibroblasts after treatment with 1mM MMS at 37°C for 30 min. NT and 0 min data replotted from (A). Repair was performed in media at 37°C. Data points and error bars represent averages and standard errors of the mean (SEM), respectively, of three independent experiments (n ≥ 3) with at least 100 comets scored for each condition in each experiment. \**p* < 0.05, Student’s *t*-test (one-tailed, paired), comparing to WT at each time point.

AAG is the major DNA glycosylase for repair of MMS-induced base lesions; specifically, AAG is responsible repairing 7MeG and 3MeA, which together comprise ∼80% of the MMS-induced base lesions. We compared cells with WT levels of AAG, cells lacking AAG (*Aag*^−/−^), and cells that overexpress the AAG through a transgenic insert (*AagTg*)(Sobol et al. 2003). For the *AagTg* cells, we observed higher levels of damage immediately after exposure to MMS (Figure 4A and B), indicative of increased initiation of BER and conversion of undetectable methylated bases into detectable BER intermediates. However, repair was not significantly different between WT and *AagTg*, possibly because Polβ is rate limiting (Horton et al. 2008; Sobol et al. 2002; Srivastava et al. 1998). We expected to see the opposite effect for AAG null cells, since there should be reduced initiation of BER, and thus reduced BER intermediates. Although there appears to be a slight suppression of BER intermediates in *Aag*^−/−^ cells just after MMS exposure, the effect was not statistically significant for any of the time points (Figure 4A and B). This could be the case if the majority of the damage observed from MMS exposure is caused by chemical induction of strand breaks, rather than enzyme-induced BER intermediates. For example, 7MeG can readily depurinate, forming AP sites, which are detected under alkaline comet assay conditions. We next analyzed the effect of knock out of Polβ. Polβ is the primary polymerase for repair synthesis following initiation of BER by AAG (Sobol et al. 1996). We found that the Polβ null cells have increased damage (Figure 4A), and that this is the case for all time points post exposure to MMS, which is consistent with inhibition of completion of the BER pathway (Figure 4B). Taken together, for both oxidative damage and alkylation damage, these studies show that the comet assay is an effective approach for monitoring completion of the BER pathway, but that the assay is less sensitive to differences in the levels of DNA glycosylases.

### Analysis of Nucleotide Excision Repair Capacity

NER is the major repair pathway for bulky DNA lesions, such as UV-induced pyrimidine dimers (Gillet and Scharer 2006; Hanawalt 2002; Marteijn et al. 2014; Scharer 2013). To remove such damage, XPF/ERCC1 and XPG cut upstream and downstream of the lesion respectively, releasing an oligonucleotide that contains the damaged base(s) (Figure 5A). NER is a highly coordinated pathway, wherein about a dozen proteins are required to assemble into a complex prior to cleavage of the DNA. As such, single-stranded intermediates are short-lived compared to BER intermediates, making it difficult to measure NER capacity. As has been shown previously (Azqueta and Collins 2013; Azqueta et al. 2014; AR Collins et al. 1997; Gedik et al. 1992), one approach for studying NER is to exploit T4 EndoV, which cleaves the DNA 5’ to the pyrimidine dimer, thus converting undetectable damage into DNA strand breaks that can be detected using the comet assay (Figure 5A).

**Figure 5.**
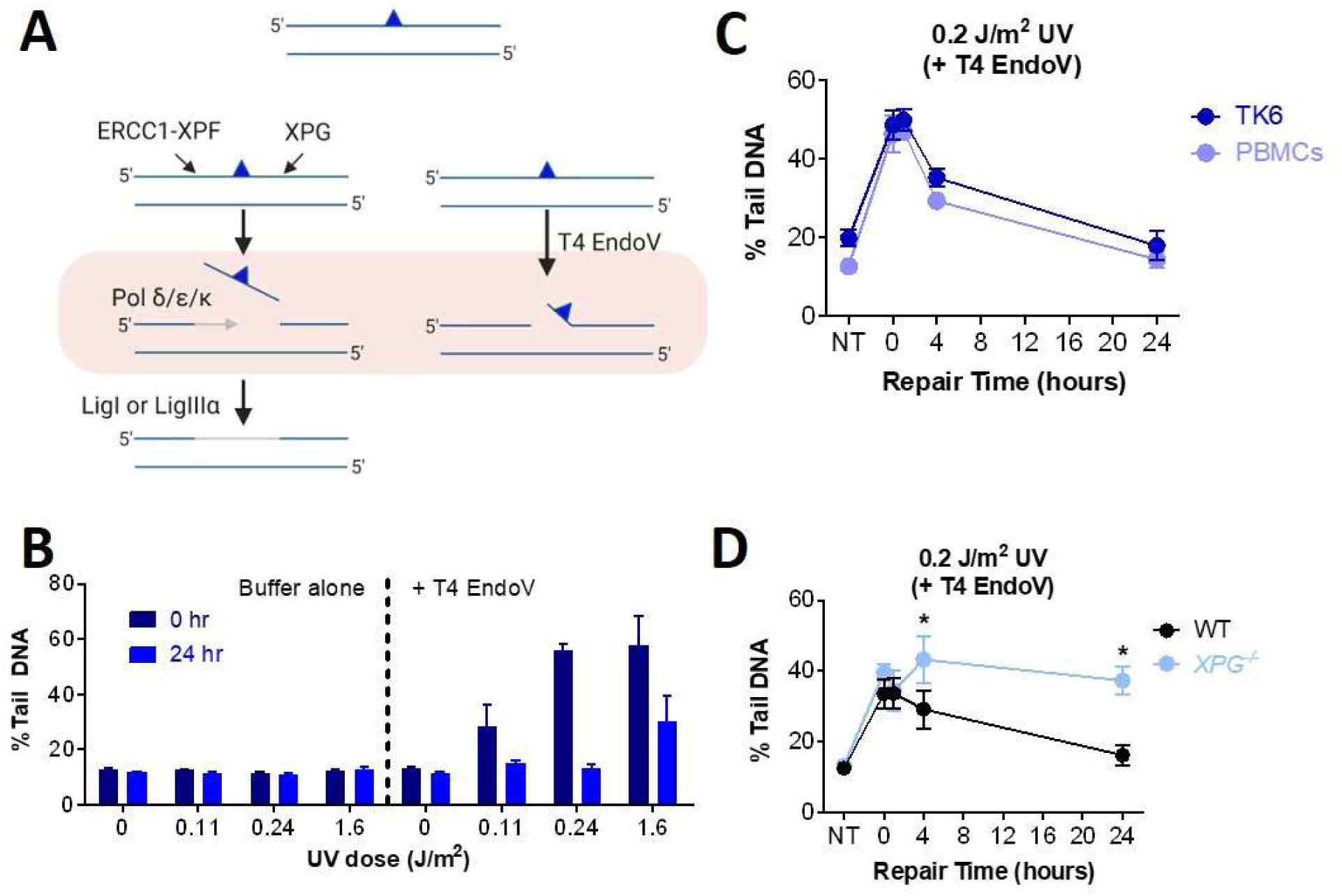
Measuring DNA NER Kinetics in Live Cells. (A) Schematic of the mammalian NER pathway with the blue triangle representing a bulky lesion. (B) UV-irradiated TK6 cells were lysed overnight, and the level of CPDs was measured by incubating the exposed nuclear DNA with T4 Endonuclease V (T4 EndoV). Buffer control is T4 EndoV reaction buffer with no enzyme added. Data points represent averages of three independent experiments. (C) TK6 and PHA-stimulated T-lymphocytes (PBMCs) after exposure to UV light. Repair was performed in media at 37°C. (D) Repair of human skin fibroblast cell lines, *XPG*^*−/−*^and WT. Data points represent averages of triplicate wells in one experiment. Error bars represent SEM. \**p* < 0.05, Student’s *t*-test (one-tailed, paired), comparing to WT at each time point.

We measured percent tail DNA for TK6 cells exposed to various doses of UV, but did not observe a dose-response, presumably because the single stranded NER intermediates are not persistent enough for detection by the comet assay (Figure 5B, left). To reveal the damage usually repaired by NER, we performed a modified comet assay, such that after exposure to UV, cells were lysed, and then subsequently incubated in buffer containing T4 EndoV (Azqueta et al. 2013; Azqueta et al. 2014; A Collins et al. 1997; Gedik et al. 1992). We observed high levels of damage just after exposure to UV (0 hr), indicative of the presence of UV dimers, which are T4 EndoV sensitive sites (Figure 5B, right). Twenty-four hours post exposure, there was a significant reduction in the levels of damage, which is consistent with repair of UV dimers by NER (Figure 5B, right).

NER activity can be highly variable in different individuals. For example, some people with Xeroderma Pigmentosum (XP) have severe defects in NER, which makes them extremely sensitive to sunlight-induced skin cancer (DiGiovanna and Kraemer 2012; Kraemer et al. 1994). NER also plays a role in removing DNA lesions induced by chemotherapeutic agents, such as cisplatin (Huang et al. 1994; Reardon et al. 1999; Wang et al. 2003; Zamble et al. 1996). Therefore, knowledge about DNA repair capacity in people has relevance both in terms of susceptibility to cancer as well as susceptibility to the toxic effects of chemotherapy. We therefore explored the utility of the CometChip for studies of NER capacity in human cells. Specifically, we monitored repair of UV-induced damage in both TK6 cells PBMCs. Using T4 EndoV, we observed a steep increase in DNA damage immediately after exposing cells to UV, followed by repair over the course of the subsequent 24 hours (Figure 5C). These results show that NER capacity is similar in transformed and primary lymphocytes, and can be effectively evaluated in primary lymphocytes using the CometChip.

To further explore the specificity of the CometChip-T4 EndoV analysis method, we studied cells with normal DNA repair and cells from an XPG-deficient patient. Since XPG is required for pre-assembly of the NER complex, cells lacking XPG do not initiate NER and strand cleavage does not occur. As a result, UV dimers are anticipated to be persistent. We tested this model by treating WT and XPG-null cells with UV, and then monitoring the disappearance of T4 EndoV sensitive sites over time (Figure 5D). Both cell types accumulated the same load of damaged DNA and in WT cells, substrates of T4 EndoV are cleared over time. In contrast, DNA damage persists in the XPG-null cells. These results show that the T4 EndoV approach specifically detects substrates of NER, providing an effective approach for testing NER status in mammalian cells.

### Measuring Non-Homologous End-Joining

Unlike the alkaline comet assay (which detects SSBs and DSBs, abasic sites and alkali sensitive sites), the neutral comet assay can specifically detect DNA double strand breaks (DSBs) (Olive et al. 1991). DSBs are repaired primarily by NHEJ and HR, and NHEJ is the primary DSB repair pathway for cells in G1 (Hinz et al. 2005; Karanam et al. 2012; Mao et al. 2008; Rothkamm et al. 2003; Takata et al. 1998). NHEJ involves recognition of the double stranded DNA ends by the Ku70/80 heterodimer, followed by recruitment of DNA protein kinase catalytic subunit (DNA-PKcs), which brings the DNA ends together in the DNA-PK complex. Ends are then processed to create blunt ends or small overhangs that are substrates for the XRCC4/LigIV/XLF complex, which ligates the DSB ends together. As examples of the utility of the CometChip for studies of NHEJ, we assayed DSB repair kinetics both for cells lacking the essential NHEJ protein DNA-PKcs and for cells treated with a chemical inhibitor of LigIV (namely, Scr7) (Srivastava et al.). We exploited the human glioma cell line, M059J, which was developed from human tumors and lacks DNA-PKcs. Its sister cell line, M059K, was derived from the same human tumor and expresses wild type DNA-PKcs. Another glioma cell line expressing wild type DNA-PKcs, LN428, was used for Scr7 treatment. After irradiation and allowing time to repair, both DNA-PKcs deficient cells and cells incubated with Scr7 demonstrated significant impairment of DSB repair (Figures 6A and B, respectively). These results are consistent with the importance of DNA-PKcs and LigIV in NHEJ repair, and agree with data indicating high potency Scr7 inhibition of LigIV (Srivastava et al. 2012). Thus, using neutral comet assay conditions on the CometChip enables studies of genetic and chemically-induced deficiencies in NHEJ.

**Figure 6.**
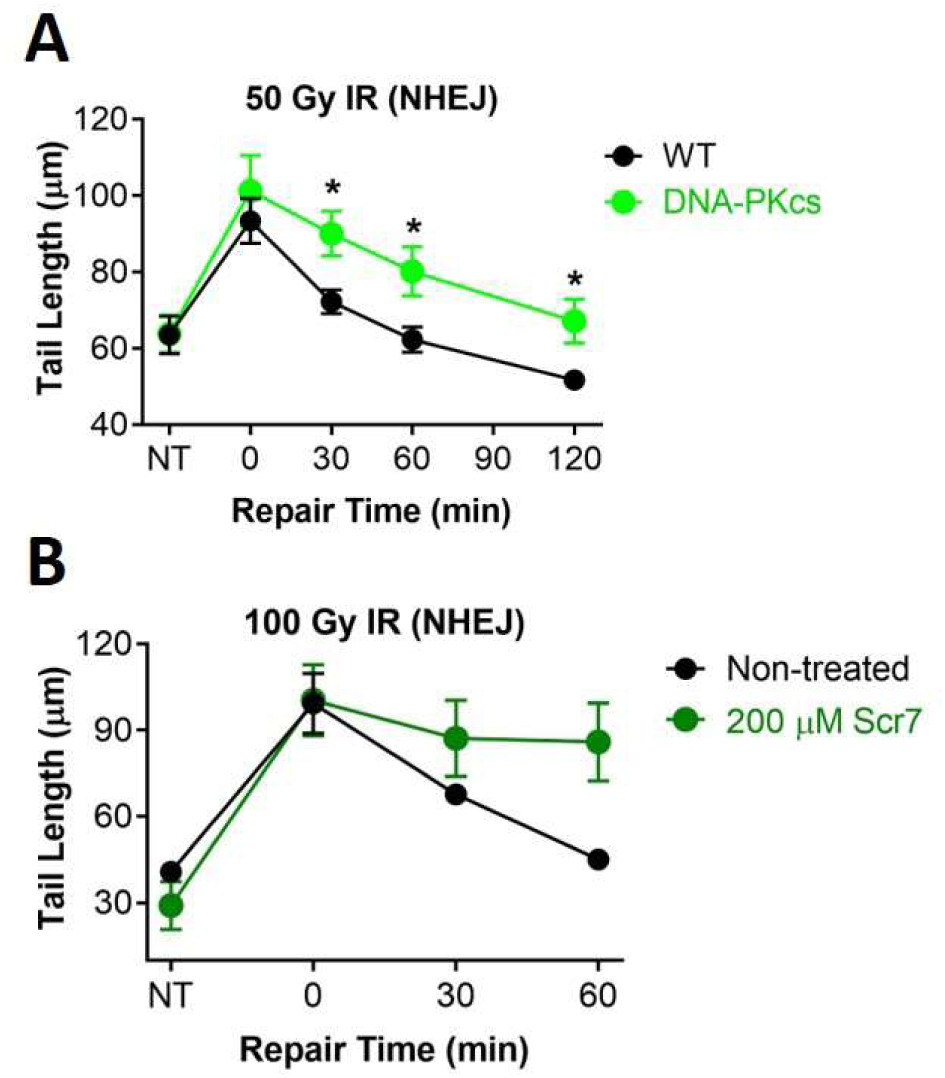
Measuring NHEJ Kinetics. (A) Repair kinetics of Glioblastoma M059K (WT) and M059J (DNA-PKcs deficient) cells after treatment with 50 Gy IR. Repair was performed in media at 37°C. (B) Repair kinetics of LN428 cells after exposure to 100 Gy IR with and without Ligase IV inhibition. LN428 cells were pre-incubated for 24 hr in media with 200 μM Scr7, a Ligase IV inhibitor. Following exposure to IR, cells were allowed to repair in media at 37°Cwith and without Scr7. Data and error bars represent averages and SEM of three independent experiments, respectively. At least 100 comets were scored for each condition in each experiment. \**p* < 0.05, Student’s *t*-test (one-tailed, paired), comparing to WT at each time point.

## DISCUSSION

Here we show that the CometChip platform enables assessment of multiple cell lines, DNA interacting agents, doses, and time points in parallel in a highly sensitive and rapid fashion. Being able to collect dense data sets with multiple time points makes it possible to assess the kinetics of DNA repair. Further, by varying either the type of DNA damaging agent and/or genetic factors, it is possible to study the repair kinetics for three major repair pathways: BER, NER, and NHEJ. Unlike the traditional assay, with the enhanced throughput of the CometChip, it is now feasible to query the impact of multiple repair deficiencies in a broad range of cell types, including primary, knock-out, knock-down, mutant, and chemically-inhibited cells. The CometChip thus has the potential to augment a broad range of applications in both the laboratory and the clinic.

Many excellent DNA repair assays exist and can provide information about a specific repair enzyme or repair pathway. However, historically there have been few assays usable for parallel analysis of both damage induction and clearance to provide detailed information on repair kinetics in living cells. Few, if any, assays have the ability to assess an agent’s potential to damage DNA and a cell’s response to the damage without prior knowledge of the agent’s mechanism of action. For example, oligo cleavage assays are good for measuring glycosylase activity but not the activity of the whole repair pathway. Other than the comet assay, currently available assays developed for BER are often limited to *in vitro* analysis of the activity of particular proteins (Li et al. 2018; Svilar et al. 2012), and thus do not reflect the repair capacity of the pathway on the whole in living cells. Available *in vivo* assays, such as unscheduled DNA synthesis (UDS) and sister chromatid exchange (SCE) are useful for NER and HR (Latimer and Kelly 2014; Wojcik et al. 2004), respectively, but they are not readily integrated into a multi-pathway platform and can be technically challenging. The development of assays for repair foci (e.g., γ-H2AX) can give real-time information on repair activity, but it is again restricted to analyses of specific repair-associated proteins and does not reflect physical DNA damage, but rather signaling events downstream of DNA damage and can be affected by multiple non-repair factors (Kinner et al. 2008; Nakamura et al. 2006). Further, γ-H2AX measures a response to DNA damage but does not directly measure DNA repair kinetics. For example, we previously assayed for DSBs in parallel with γ-H2AX and found that physical DSBs were quickly cleared, while a γ-H2AX signal persisted for more than 6 hours afterwards (Weingeist et al. 2013). Recently, the development of the FM-HCR multiplexed platform provides for the first time the potential to assess multiple repair pathway capacities in parallel by inserting exogenously damaged DNA into living cells (Nagel et al. 2014b). It is noteworthy that DNA damaging agents generally induce a range of DNA lesions that are repaired by multiple DNA repair pathways, and thus DNA repair kinetics may vary by chemical agent. By analyzing physical strand breaks, the comet assay offers a complementary approach to FM-HCR in its ability to measure agent-specific DNA repair kinetics in a natural chromatin context. Importantly, the CometChip platform incorporates well-established and validated comet protocols (Collins 2004; Olive and Banath 2006; Sykora et al. 2018; Tice and Strauss 1995). In combination with the micro-patterning technology that is the basis for the CometChip, the studies presented here show that this is a fast, simple, and reliable method to study the global repair response. Both FM-HCR and the CometChip have the strengths of versatility and adaptability to measure multiple repair pathways in living cells.

In conclusion, we have demonstrated that the CometChip is an effective platform for assessing repair kinetics across several classes of DNA damage addressed by three major DNA repair pathways in living cells, namely BER, NER and NHEJ.As such, this platform opens doors to a deeper mechanistic understanding of DNA repair and discovery of multi-functionality of key DNA repair proteins. Clinically, the CometChip platform can also be used to understand how candidate drugs work mechanistically, to study on- and off-target effects, and to optimize treatment regimens. Finally, for public health, the CometChip can be a potential biomonitoring device for studying effects of environmental genotoxins and for evaluating variation in repair responses among people and among populations. The versatility, simplicity, high throughput capacity, and low cost of the CometChip together make this platform valuable for basic, clinical and public health research.

## ACKNOWLEDGEMENTS

We thank Dr. Zachary Nagel for his critical comments. We thank Dr. Samuel H. Wilson for all mouse embryonic fibroblast cell lines. We thank Dr. David K. Wood for his work on developing the CometChip and the analysis program.

## AUTHOR CONTRIBUTIONS

B.P.E. conceptualized the study; J.G., L.P.N., I.J.T., E.T., D.N.C., J.L.F., D.M.W., P.M. performed the CometChip experiments. J.G., L.P.N., I.J.T., E.T., D.N.C., J.L.F., D.M.W., S.K. analyzed the data. J.G., B.P.E., S.K., J.E.K. drafted the manuscript. B.P.E., L.D.S., R.W.S., and S.R.F. acquired funding.

